# Ancient cyanobacterial proto-circadian clock adapted to short day-night cycles ∼ 0.95 billion years ago

**DOI:** 10.1101/2024.04.30.591965

**Authors:** Silin Li, Zengxuan Zhou, Yufeng Wan, Xudong Jia, Peiliang Wang, Yu Wang, Shuqi Dong, Jun He, Xujing Wang, Ximing Qin, Qiguang Xie, Xiaodong Xu, Yuwei Zhao, Dan Liang, Peng Zhang, Qinfen Zhang, Jinhu Guo

## Abstract

In the early history, the Earth rotation was much faster; however, how ancient organisms adapted to the day-night cycles of that time remains unclear. We resurrected the ancestor KaiABC (anKaiABC) genes circa 0.95 billion years (Ga) ago when the daily-night cycling period was ∼ 18 h. Compared with its contemporary counterpart KaiC, the anKaiC protein shows subtle structural differences, and the activities of kinase, phosphatase activities, and adenosine triphosphatase (ATPase) of anKaiC are lower. The anKaiAB proteins are less effective in regulating KaiC/anKaiC phosphorylation status. The anKaiABC system does not sustain an endogenous oscillation but it can be entrained by an 18-h light/dark cycle. The strain expressing *anKaiABC* shows better adaptation under 9-h light/9-h dark cycles (LD9:9) which mimic the 18-h day-night cycles. These findings suggest that the ancient cyanobacterial proto-circadian system may not be endogenous, but it conferred the capability to adapt to daily cycles ∼ 0.95 Ga ago.

## INTRODUCTION

The Earth rotates on its axis, and as a consequence, many environmental factors, e.g., light, temperature, humidity, and radiation, periodically alternate. Across kingdoms, the circadian system has evolved to fit with the 24-h light/dark cycle owing to the rotation of the Earth. Dominant eukaryotic organisms possess sustainable circadian clock systems with periods of circa 24 h, which endow them with the capability to synchronize their physiology and behaviour to daily environmental alternations. ^1,2^ Circadian rhythms are also present in some prokaryotic species, e.g., cyanobacteria and nonphotosynthetic *Bacillus subtilis*. ^3–5^ It has been demonstrated in cyanobacteria and some other species, that a circadian clock with a period close to the environmental cycling alternation enhances fitness or adaptation.^6,7,8–11^

Cyanobacteria are regarded as the earliest organisms to have emerged up to 3.5 Ga ago^12^. Circadian rhythms in multiple physiological variables, e.g., nitrogen fixation and photosynthesis, have been found in several cyanobacteria species including *Synechococcus* RF-1, Miami BG 43511/43522, WH7803 and *Synechococcus elongatus* PCC 7942. *S. elongatus* PCC 7942 has an endogenous circadian system, which consists of KaiABC proteins.^13,14^ KaiC is both an autokinase and an autophosphatase that contains two phosphorylation sites: S431 and T432. ^15^ KaiC phosphorylation is correlated with the dual adenosine triphosphatase domains located in the CI (1-268 aa) and CII (269-519 aa) regions, respectively. ^16^ KaiA binds to the KaiC hexamer and promotes KaiC phosphorylation while KaiB binds to KaiC which sequesters KaiA from KaiC A-loop (488-497 aa). ^17,18^ Interestingly, even in vitro, the KaiABC proteins display intrinsic oscillation of KaiC phosphorylation and desphosphorylation. ^19^ The evolutionary history of some cyanobacterial circadian genes, e.g., *kaiC*, *sasA*, *cikA* and *cpmA*, can be dated back to about 3.5 Ga ago. It has been proposed that KaiA might have appeared in cyanobacteria genomes 1.6 Ga – 1.4 Ga ago or even earlier. ^20,21^ Some modern cyanobacteria, e.g., *Gloeobacter violaceus* PCC 7421, *Prochlorococcus sp*. strain MED4, *Rhodopseudomonas palustris*, and *Rhodobacter spheroides*, contain no *KaiA*, suggesting that they have no endogenous self-sustaining circadian clocks. ^22,23–25^

The rotation of Earth is slowing mainly due to gravitational pull by the moon, and the rotation period was much faster in ancient times, as evidenced by astronomical measurements and palaeontological data. ^1–3^ The ancient environment on Earth dramatically differed from the present environment, including in terms of the period of daily light/dark cycling, and was much shorter due to the Earth’s faster rotation. For instance, 0.9 Ga ago, the day length was ∼ 18 h. ^29^ Analysis of mollusc fossils suggests that the ancient animals had shorter days than they do today. ^30^ However, to date, there is very limited knowledge regarding this phenomenon at the molecular level.

It is very intriguing but challenging to investigate the characteristics of ancient circadian systems and their contribution to adaptation to the ancient environment. In this study, we predicted and reconstructed an ancient cyanobacterial proto-circadian clock system ∼ 0.95 Ga ago, and the functional characteristics and adaptation to the daily cycles of the temporal environment were analysed.

## RESULTS

### Phylogenetic analysis of cyanobacterial KaiABC proteins

Ancient proteins can be reconstructed and their biochemical properties and functions can be assayed based on phylogenetic analysis of their evolutionally contemporary descendents. ^31,32^ During the early history of the Earth, the planet rotated much faster than now which means that the period of daily light/dark cycling was much shorter. ^29,30^ To investigate whether ancient circadian clock was aligned with the ancient light/dark cycling environment, we searched the KaiABC sequences of all available extant cyanobacterial species and performed a phylogenetic analysis. The candidate ancestral sequences were then reconstructed at nodes throughout the bacterial subtree, and the results indicated that the nodes of ancestral KaiABC sequences simultaneously occurred ∼ 0.95 Ga ago based on the available KaiABC sequences from modern cyanobacterial species (Figure 1A-C). The divergence time was deduced according to the 16S rRNA gene information. ^33^ The overall sequences of critical motifs in anKaiABC proteins were highly conserved with those in KaiABC, especially in KaiC despite the dispersed amino acid residues (Figure S1A-C). Based on these phylogenetic results and sequence alignments, we synthesized the ancestral genes that express these deduced ancestral KaiABC proteins (anKaiABC) from ∼ 0.95 Ga ago.

**Figure 1.**
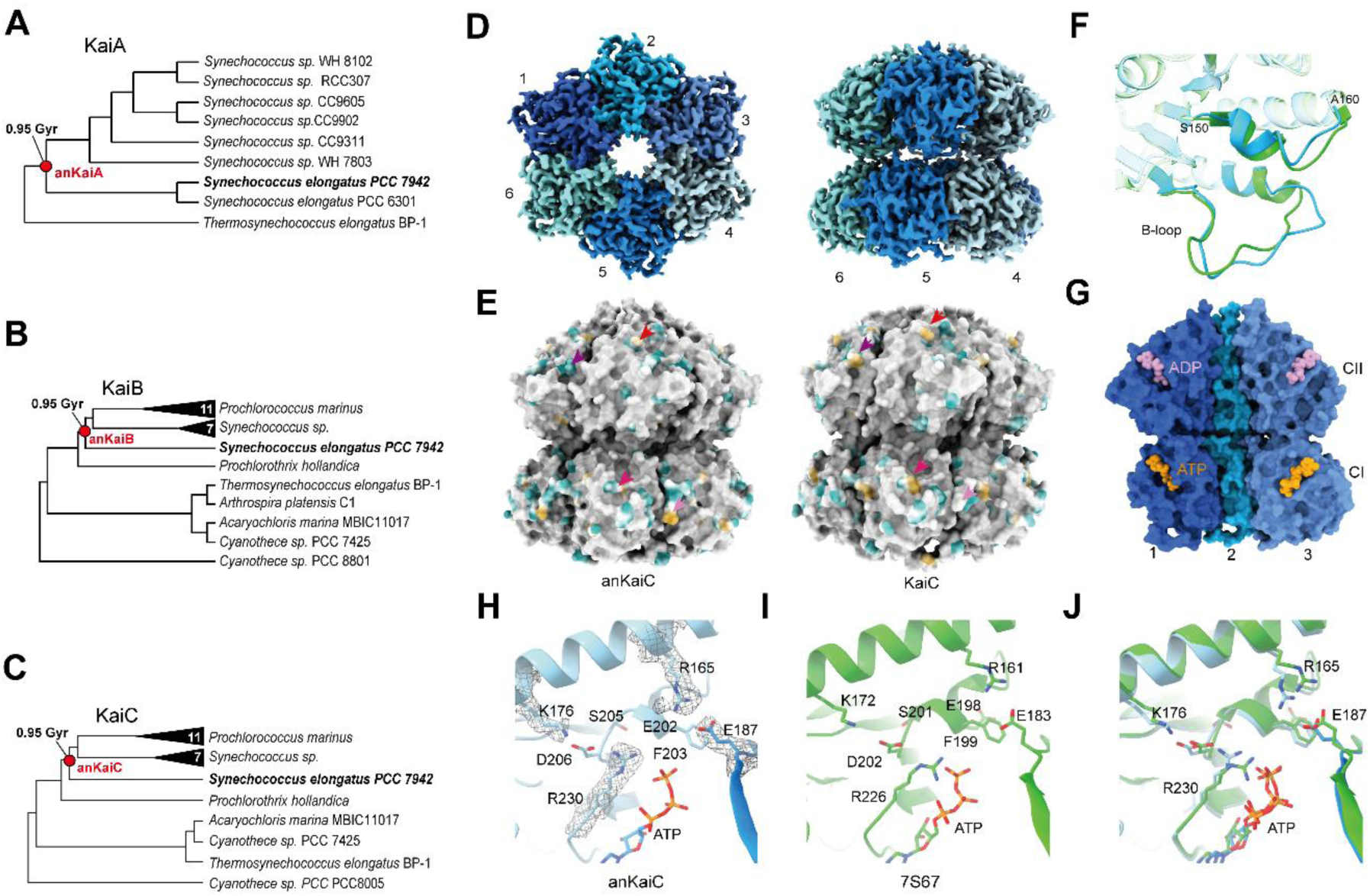
Analysis of ancient KaiABC (anKaiABC) proteins. (A-C) Unrooted universal phylogenetic trees used to reconstruct ancestral KaiABC proteins. The branching time of ∼0.95 Ga. d-f: *In vitro* assays to characterize the anKaiABC proteins. Structural analysis of anKaiC. (D) Density map of the anKaiC hexamer viewed from the top (left panel) and side (right panel). Monomers are labelled. The six anKaiC monomers are marked by numbers and coloured with different colours. (E) Structural differences between anKaiC (8JON) and KaiC WT (PDB: 7S67). The surface (outside view) with identical amino acid residues is coloured in grey, the different amino acids between anKaiC and KaiC with hydrophilic residues are shown in cyan, and the hydrophobic residues are shown in goldenrod. The arrows in different colours denote the corresponding areas with different hydrophobicities due to different amino acid residues. (F) Alignment of models of anKaiC and KaiC (PDB:7S67) indicates both B-loop and the loop (S150-A160) of anKaic cannot align well with that of KaiC. (G) Interface representation of anKaiC (including monomers 1, 2 and 3) viewed from inside, with ATP nucleotides displayed in orange and ADP in pink. (H,I) The amino acids of anKaiC (H) and Kai C (I) surrounding the ATP. (A) (J) Alignment of models of anKaiC and KaiC (PDB:7S67) indicates several side chains of the amino acids around the ATP have subtle difference in location and orientation. See Figures S1-3.

### Cryo-EM analysis of the anKaiC structure

anKaiABC proteins were prokaryotically expressed and purified (Figure S2). We analysed the structure by cryo-EM single particle reconstruction, and obtained the structure of anKaiC with a resolution of 2.5 Å (Figure S3A-E; Table S1). anKaiC can form a conserved homohexamer, which bears a highly conserved structure with KaiC (Figure 1D) (PDB:7S67).

**Figure 2.**
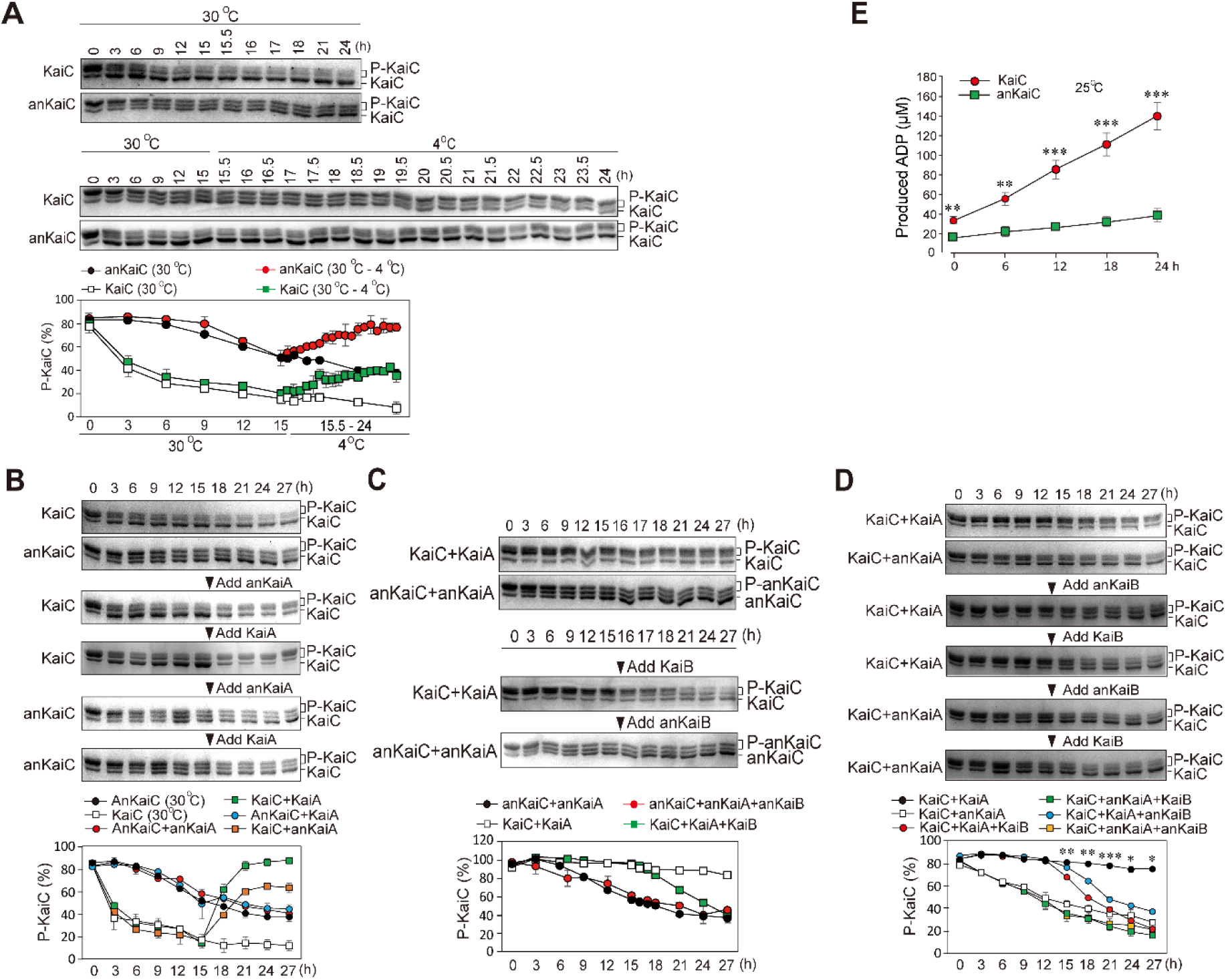
Functions of ancient KaiAB in the regulation of KaiC/anKaiC phosphorylation. (A) Results of temperature switch experiments to compare the autokinase activities between anKaiC and KaiC. (B) Function of anKaiA in promoting anKaiC/KaiC phosphorylation. (C) Function of anKaiB in promoting anKaiC/KaiC dephosphorylation. (D) Reciprocal assays of anKaiB/KaiB in promoting anKaiC/KaiC dephosphorylation. The results were visualized with Coomassie brilliant blue staining. n=3. Asterisks denote significance between the combination of KaiA+KaiB+KaiC and KaiA+KaiB+anKaiC. (E) ATPase assay results of KaiC and anKaiC. The ADP concentration in the reaction mixture was measured by LC–MS as described. Data are means ± s.d. (A-D) and s.e.m (E). n=3.

^17^ Dispersed differences in the surface hydrophobicity and structure between KaiC and anKaiC were observed (Figure 1E). The A-loop is the binding site for KaiA to facilitate KaiC phosphorylation. ^34^ The sequences of the detectable part of anKaiC A-loop (R492 - T499, S-shaped loop^29^) are almost identical to those in KaiC, and they have a very similar structure in this region (Figure S3F). However, whether the structure of the rest C-terminal part (R493 - S524) of A-loop is similar remains unclear because this region is so flexible that the structure cannot be resolved.

In KaiC the binding site for KaiB is called B-loop (116–123 aa), and the KaiB-KaiC interaction also leads to disassociation of KaiA from KaiC A-loop and binding with KaiB. ^35^ This way KaiA is sequestered from KaiC which causes its dephosphorylation. ^34^ Differences in the B-loop (A112-D126) and another spatial loop (S150-A160) close to the B-loop were observed although its function remains elusive (Figure 1F). anKaiC harbours two clefts for the binding of ATP molecules in its CI (KaiC-CI) and CII domains like KaiC ^15^ (Figure 1G). The KaiC-CI region exhibits extremely weak but stable ATPase activity. ^36^ At the structural level, the extremely slow ATPase reaction of Kai-CI region is due to the sequestration of lytic water molecule from the γ-phosphate of ATP and cis-to-trans isomerization of the peptide bond between residues D145 and S146. ^37^ We compared the structure of ATP-binding sites in the CI region between anKaiC and a recently published structure of full-length KaiC also derived from cryo-EM analysis (PDB:7S67). ^17^ Although there is marginal difference between the inside surface hydrophobicity between anKaiC and KaiC (Figure S3G), the position and orientation of side chains amino acid residues (R165, E187, D206, and R230) which are surrounding the ATP molecule, showed subtle differences compared to the features of modern KaiC. Moreover, subtle differences in the position and orientation of ATP molecule were also observed (Figure 1H-J). These structural discriminations may cause functional differences in anKaiC in the catalytic capabilities as an ATPase, kinase or phosphatase.

### Characterization of ancient KaiABC protein functions

Cyanobacterial KaiC is both an autokinase and a phosphatase. ^38^ At high temperature, KaiC undergoes dephosphorylation. ^14^ Similarly, anKaiC also showed a decrease in phosphorylation at 30°C but the rate was much slower (Figure 2A). Shifting to a low temperature (4°C) after exposure to 30°C for 15 h led to a significant increase in the phosphorylation level of both KaiC and anKaiC, although the increase in anKaiC phosphorylation was significantly lower than that of KaiC (Figure 2A). These data suggest that anKaiC has autokinase and phosphatase activities.

To assess the activity of anKaiA in promoting KaiC/anKaiC phosphorylation, KaiA and anKaiA were mixed with KaiC/anKaiC and the results showed that anKaiA failed to promote anKaiC phosphorylation in contrast to the significant effects of KaiA on KaiC (Figure 2B). Next, we assessed the function of anKaiB in dephosphorylating anKaiC, and the results showed that compared to KaiB, anKaiB showed an undetectable function in repressing anKaiC dephosphorylation. In the combination of KaiC and KaiA (Figure 2C), the addition of anKaiB led to decreased KaiC phosphorylation, but its capability to facilitate KaiC dephosphorylation was lower than that of KaiB (Figure 2D). Together, these data suggest that ancient cyanobacterial clock components are less efficient activities in modulating KaiC phosphorylation and dephosphorylation than are their contemporary counterparts. We next compared the ATPase activities (ATP hydrolysis) between anKaiC and KaiC. The production of ADP was monitored by liquid chromatography–mass spectrometry (LC–MS). ^29^ The ATPase activity of anKaiC was significantly lower than that of KaiC (Figure 2E).

### Characterization of the ancient cyanobacterial clock system rhythmicity

The characteristics of anKaiABC proteins suggest that the circadian clock function of anKaiABC may differ from the modern cyanobacterial circadian clock system. Since KaiABC produces sustainable KaiC phosphorylation rhythms in vitro, ^17^ we conducted an in vitro oscillation assay and mixed KaiABC and anKaiABC proteins in multiple combinations to assess their circadian functions. As control, KaiABC showed robust circadian rhythms with a period of ∼ 22 h. In contrast, all of the other combinations showed no detectable rhythms, although anKaiB showed a promotion of KaiC phosphorylation at 30-40 h together with KaiA (Figure 3A,B; Figure S4), suggesting that anKaiB may possess a function closer to that of its modern counterpart.

**Figure 3.**
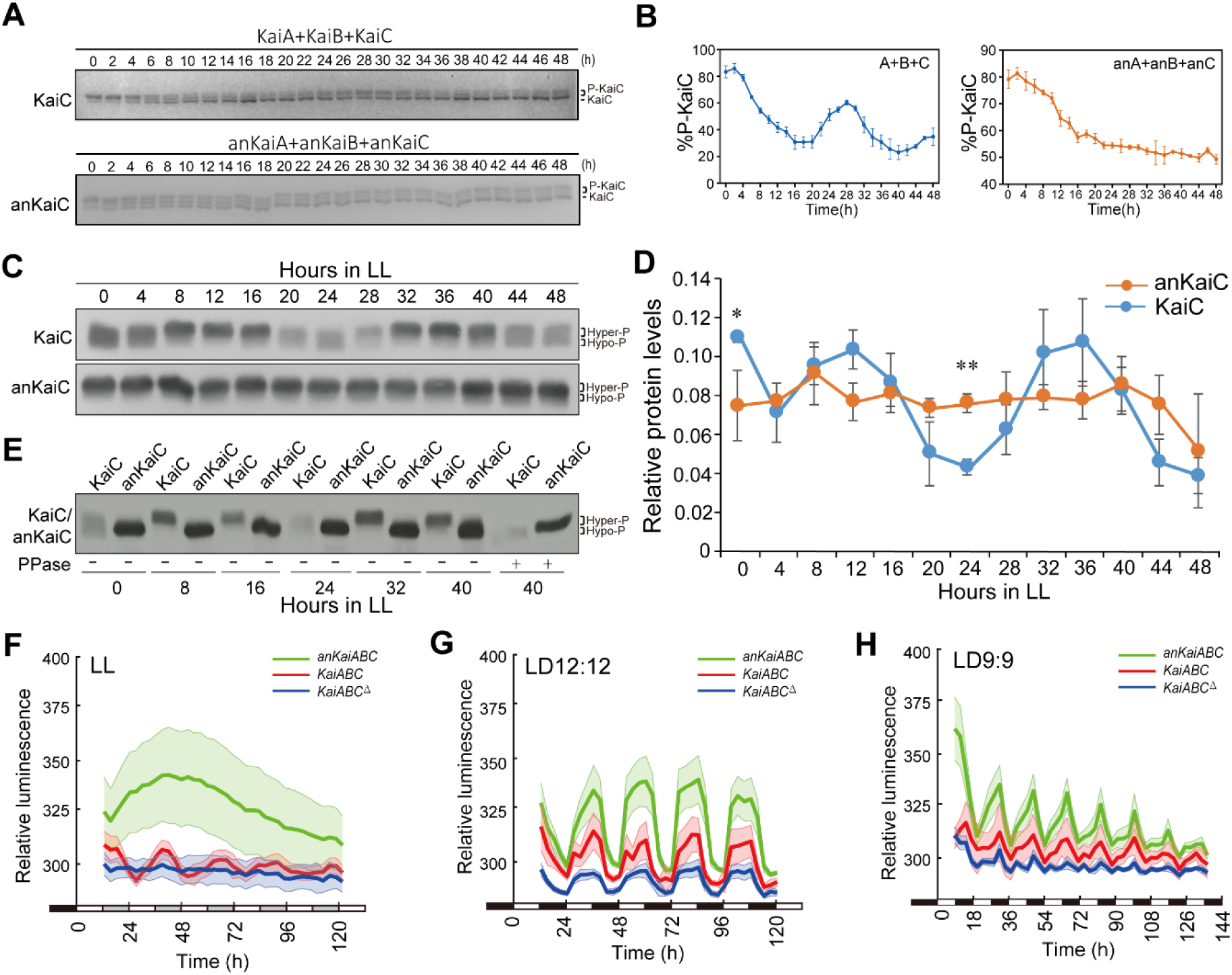
Analysis of the circadian rhythmicity of the ancient circadian system. (A,B) Representative in vitro phosphorylation of anKaiC/KaiC proteins. “P-KaiC” denotes phosphorylated KaiC proteins and “KaiC” denotes unphosphorylated KaiC proteins. The percentage of hyperphosphorylated KaiC was calculated. Representative results of Coomassie brilliant blue staining and curves of independent triplicates are shown. (C,D) Representative Western blot results of KaiC/anKaiC in LL (C) and the statistical results (D). Hyper-P denotes hyperphosphorylated proteins and hypo-P denotes hypophosphorylated proteins. Asterisks denote significance between KaiC and anKaiC. Data are means ± s.e. n=3. (E) Comparison of KaiC/anKaiC phosphorylation in LL. n=3. (F-H) Bioluminescence rhythms of *KaiABC*, *anKaiABC* and *KaiABC*^Δ^ strains. n>50. Data are means ± s.d. (F-H). See Figure S4.

**Figure 4.**
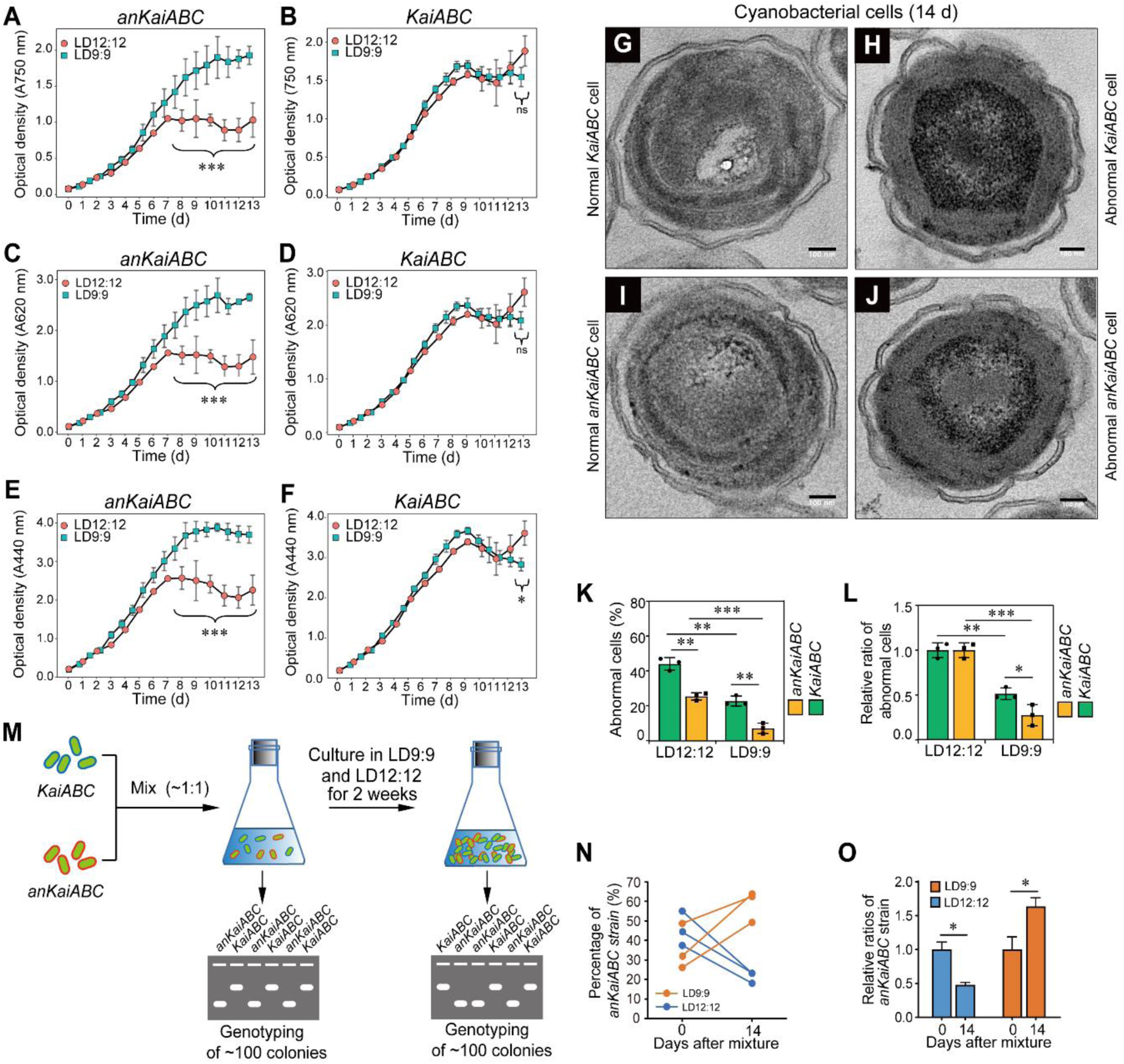
Contribution of ancient cyanobacterial circadian clock system to growth and adaptation. (A,B) Growth curves of the indicated strains in LD9:9 and LD12:12 conditions. n=3. (C,D) Phycobilin content measured by light absorption at 620 nm. n=3. (E,F) Content of chlorophyll a measured by light absorption at 440 nm. n=3. (G-J) Observation of cells by transmission electron microscopy. Bars represent 100 nm. Triplicates of samples were observed and each group comprised ∼ 100 cells. (K,L) Statistics of the results from (G-J). n=100. Scales denote 100 nm. (O) Illustration of the competition experiment. Two initial strains of interest are mixed (∼1:1) and after approximately 15 days single colonies derived from the culture are subjected to genotyping to determine the ratio of the two strains. The initial strains are denoted in different colours. (M) Schematic of the competition experiment. PCR of approximately 100 colonies was conducted to screen and confirm their genotypes. (N,O) Competition results between *KaiABC* and *anKaiABC* strains. Triplicate results are shown, and in each experiment, *n* > 100. Data are means ± s.d. (A-F) and means ± s.e.m. (I,K). See Figure S5.

To further validate whether the anKaiABC system can drive endogenous circadian rhythmicity, we transformed the *anKaiABC* genes into the *KaiABC* null strain (*KaiABC*^Δ^) of *Synechococcus sp*. PCC7942 with luciferase reporter driven by the *KaiBC* promoter (P*kaiBC*::*luxAB*). ^13^ We then conducted western blot experiments to compare the expression and phosphorylation of KaiC/anKaiC between *KaiABC* and *anKaiABC* strains under different light/dark conditions. In the constant light (LL) condition, robust circadian rhythms of abundance and phosphorylation were observed in the *KaiABC* strain but not the *anKaiABC* strain (Figure 3C-E).

Under LL conditions, an in vivo bioluminescence assay revealed that the *KaiABC* strain, but not the *KaiABC*^Δ^ and *anKaiABC* strains, showed overt circadian rhythms of bioluminescence (Figure 3F). In contrast, all three strains showed overt bioluminescence rhythms in both LD12:12 (Figure 3G), and *KaiABC* and *anKaiABC* strains but not *KaiABC*^Δ^ showed overt bioluminescence rhythms in LD9:9 which mimics the 18-h daily period approximately 9.5 Ga ago (Figure 3H). These data suggest that an ancient cyanobacteria circadian system can be entrained by environmental cues (light/dark cycles) despite the absence of endogenous rhythmicity.

### Adaptation of the ancient clock system to 18-h light/dark cycles

The strains were monitored at 750 nm in LD9:9 and LD12:12 to record the growth rate. For the *anKaiABC* strain, its growth rate before 6 d was similar in LD12:12 and LD 9:9, but after 6 d the growth rate in LD9:9 was dramatically higher than that under LD12:12. In contrast, the growth rates of *KaiABC* strain were comparable, although a slight higher but non-significant growth between LD12:12 and LD9:9 was observed on 13 d (Figure 4A,B). The absorption values at 620 nm and 440 nm wavelengths of light exposure were also measured to quantify the relative density of the two photosynthetic pigments, phycobilin and chlorophyll a, respectively. ^39^ The results showed that in the *anKaiABC* strain, similar to the growth curves, both the contents of phycocyanobilin and chlorophyll a were significantly higher in LD9:9 compared to LD12:12. However, the KaiABC strain showed comparable levels of phycobilin between LD12:12 and LD9:9, and a slightly higher level of chlorophyll in LD12:12 than that in LD9:9 on 13 d (Figure 4C-F). Interestingly, the *KaiABC^Δ^* strain showed different curves of growth and photosynthetic pigment contents from those in the *anKaiABC* strain (Figure S5A-F).

The transmission electron microscopy (TEM) results revealed that some cyanobacterial cells showed abnormal phenotypes of crenated cell walls and dissembled phycobilisomes (Figure 4G-J). The ratios of the abnormal cells of both the *anKaiABC* and control strains were significantly higher at 14 d than those at 1 d, demonstrating the exhaustion of nutrition due to excessive growth (Figure 4K). Interestingly, the increase in the ratio of abnormal cells was significantly lower for the *anKaiABC* strain than for the control in LD9:9 (Figure 4L).

Moreover, we conducted a competition experiment with these two strains (Figure 4M) by following a previous protocol. ^6^ The ratios of each strain were determined by genotyping with primers targeting different sequences (Table S2). As illustrated in Figure 4N,O, *KaiABC* grew better during 14 d of competition in the 24-h cycle, but the *anKaiABC* outcompeted *KaiABC* in the 18-h cycle. Together with the results from growth experiments, these data demonstrate that compared to other strains, the *anKaiABC* strain showed a preferable growth in LD9:9.

## DISCUSSION

The palaeontologic evidence from the daily or monthly growth rates of corals and shells demonstrates that the rotation of Earth is slowing and the day length is increasing. ^40^ However, fossils cannot tell the dynamic story of the circadian clock at the molecular level. The resurrection of the ancient cyanobacterial clock system may provide an evolutionary tool for understanding the features and function of the ancient circadian system. ^32^

The KaiABC system is essential for cyanobacteria to generate circadian rhythms. The marine genus *Prochlorococcus* can be strongly synchronized by the daily environmental cycles. ^41,42^ *Prochlorococcus* contains KaiBC but not KaiA, which explains the absence of endogenous circadian rhythmicity. In contrast, *Synechococcussp*. WH8102 comprises intact clock components and it displays endogenous circadian rhythms. ^22^

It is possible to investigate the shift in function and/or structure of ancestor proteins by reconstructing a protein family phylogeny which represents a discrete interval of evolutionary time. ^31^ For instance, Gaucher et al resurrected the elongation factors of the Tu family (EF-Tu) according to the ancient nodes in the bacterial evolutionary tree, and found that the ancient EF-Tu proteins display optimal activities at temperature of 55°C–65 °C. ^32^ The mtDNA sequence of a single African female ancestor can be traced back from the contemporary populations. ^42^ In this study, we reconstituted the ancient cyanobacterial circadian clock system of KaiABC proteins, and entrainment in LD12:12 and LD9:9 implies that the anKaiABC proteins can be integrated with circadian input and output pathways in contemporary cyanobacterial cells (Figure 3G,H).

KaiC is the core component in the KaiABC system, and it is capable of phosphorylating and dephosphorylating itself. ^43–45^ Likewise, anKaiC also forms hexamers, as revealed by cryo-EM (Figure 1D). In vitro biochemical assays revealed that anKaiA can effectively promote the phosphorylation of KaiC despite its activity being lower than that of KaiA. In contrast, anKaiA showed no function in phosphorylating anKaiC (Figure 2B). Some species with primitive circadian system lacking kaiA cannot generate self-sustainable oscillation, suggesting the critical role of KaiA in maintaining the function of cyanobacterial circadian clock. ^10,32,46^ anKaiB has a partial function in promoting the dephosphorylation of KaiC but not anKaiC (Figure 2D). Moreover, anKaiC showed dramatically lower activity in ATP hydrolysis (Figure 2E). Consistently, the structural difference in B-loop region may account for the lower phosphatase activity of anKaiC (Figure 1F); the subtly different structure in ATP-binding region of anKaiC may explain the lower ATPase activity of anKaiC (Figure 1H-J). The C-terminal part of A-loop in anKaiC has several amino acid residues different from those in KaiC (Figure S1C; Figure S3F), suggesting this region may have a different structure from anKaiC which accounts for the lower level of anKaiC phosphorylation through affecting KaiB-KaiC interaction.

In addition, the combination of anKaiA-anKaiB-anKaiC showed no circadian rhythms in KaiC phosphorylation and the phosphorylation level of KaiC was constantly low (Figure 3A,B). Consistently, in the 24-h cycles, the *KaiABC* strain also displayed circadian rhythms at the KaiC level, while the anKaiABC strain was arrhythmic (Figure 3C-E). Together, these data demonstrate that compared to the modern KaiABC, the ancient KaiABC system already had a partial circadian clock function; however, it was still too evolutionarily immature to maintain a self-sustainable circadian oscillation. A sustainable circadian system may have developed some time after 9.5 Ga ago in cyanobacteria.

It has been proposed that the ATPase activity of KaiC correlates with the circadian period length, ^37^ which suggests that anKaiC should be more active in catalysis of ATP hydrolysis. However, the present study shows that the ATPase activity of anKaiC is much lower than contemporary KaiC (Figure 2E). This inconsistency could be explained as that the correlation may only apply to the KaiABC system which already possesses self-sustainability, but not the developmentally and functionally immature ancestor.

Despite the lack of an endogenous circadian clock in the *anKaiABC* strain, we showed that the bioluminescence rhythms of this strain can be entrained in the LD12:12 and LD9:9 conditions (Figure 3G-H), suggesting that the ancient anKaiABC system may constitute an hourglass-type clock. Supportively, it has been demonstrated that in different bacteria, an hourglass-type clock can also coordinate physiological processes to adapt to the cycling environment. ^10,32,46,47^ *Cyanobacteria* expressing ancient circadian genes showed significantly increased growth rate and higher content of photosynthetic pigments after exhaustive growth in LD9:9 for 7 d (Figure 4A-F), and competitive strength in LD9:9 (Figure 4M-O). Furthermore, the *anKaiABC* strain showed no overt entrained bioluminescence rhythmicity in LD9:9, and exhibited different growth and photosynthetic pigment levels compared with the *KaiABC*^Δ^ strain (Figure 3H; Figure S5A-F), suggesting that the ancient proto-circadian system already possessed certain function in modulating diurnal rhythms and fitness to the environment despite its lack of self-sustainability. As the content of atmospheric O2 ∼0.95 Ga ago was less than 1/10 of current levels, it is likely that the ancestral circadian system may confer cyanobacteria adaptability, enabling them to grow at a higher density under the low oxygen environment. ^48^

Together, these data suggest that the ancient proto-circadian clock system is beneficial for cyanobacteria survival in the 18-h daily cycles ∼ 0.95 Ga ago, and that the ancient cyanobacterial circadian components already had certain functions in regulating growth or metabolism despite the lack of endogenous oscillation.

## Data availability

The cryo-EM maps of anKaiC were deposited in the Electron Microscopy Data Bank (EMDB) under the accession code EMD-36461. The atomic coordinates were deposited in the RCSB Protein Data Bank (PDB) under the accession code 8JON.

## ACKNOWLEDGEMENTS

We thank Y. Xu and C.H. Johnson (Vanderbilt University) for providing strains and technical help; Y. Liu (UT Southwestern Medical Center) and Y. Li (Hongkong University) for helpful discussion; D. Chen and W. Zhuo (Sun Yat-sen University) for technical support. Research reported in this publication was supported by funding from the National Key R&D Program of China 2022YFA1604504 (to J.G.), the Lin Gang Laboratory & National Key Laboratory of Human Factors Engineering Joint Grant LG-TKN-202203-01 (to J.G.), the Experiments for Space Exploration Program and the Qian Xuesen Laboratory, China Academy of Space Technology TKTSPY-2020-04-21 (to J.G.), the Open Fund of the National Key Laboratory of Human Factors Engineering in the Astronaut Center of China SYFD062008K (to J.G.) and the National Natural Science Foundation of China 32171162 (to J.G.).

## AUTHOR CONTRIBUTIONS

J.G. conceptualised the project; Funding acquisition: J.G.; S.L., Y.W., P.W., D.L. and P.Z. predicted the ancient KaiABC sequences. S.L., Z.Z., P.W., Y.W., X.W., X.Q., Q.X., X.X. and Y.Z. performed the experiments. S.L., X.J., S.D., J.H. and Q.Z. conducted the cryo-EM experiments; J.G. supervised the project. S.L., J.G., X.J, and Q.Z. wrote the paper.

### Additional information

**Extended data** is available for this paper including Extended Figs. 1-4 and Extended Tables 1,2.

**Supplementary information** The online version contains uncropped western blots and PCR electrophoresis results.

## STAR★METHODS

## KEY RESOURCES TABLE

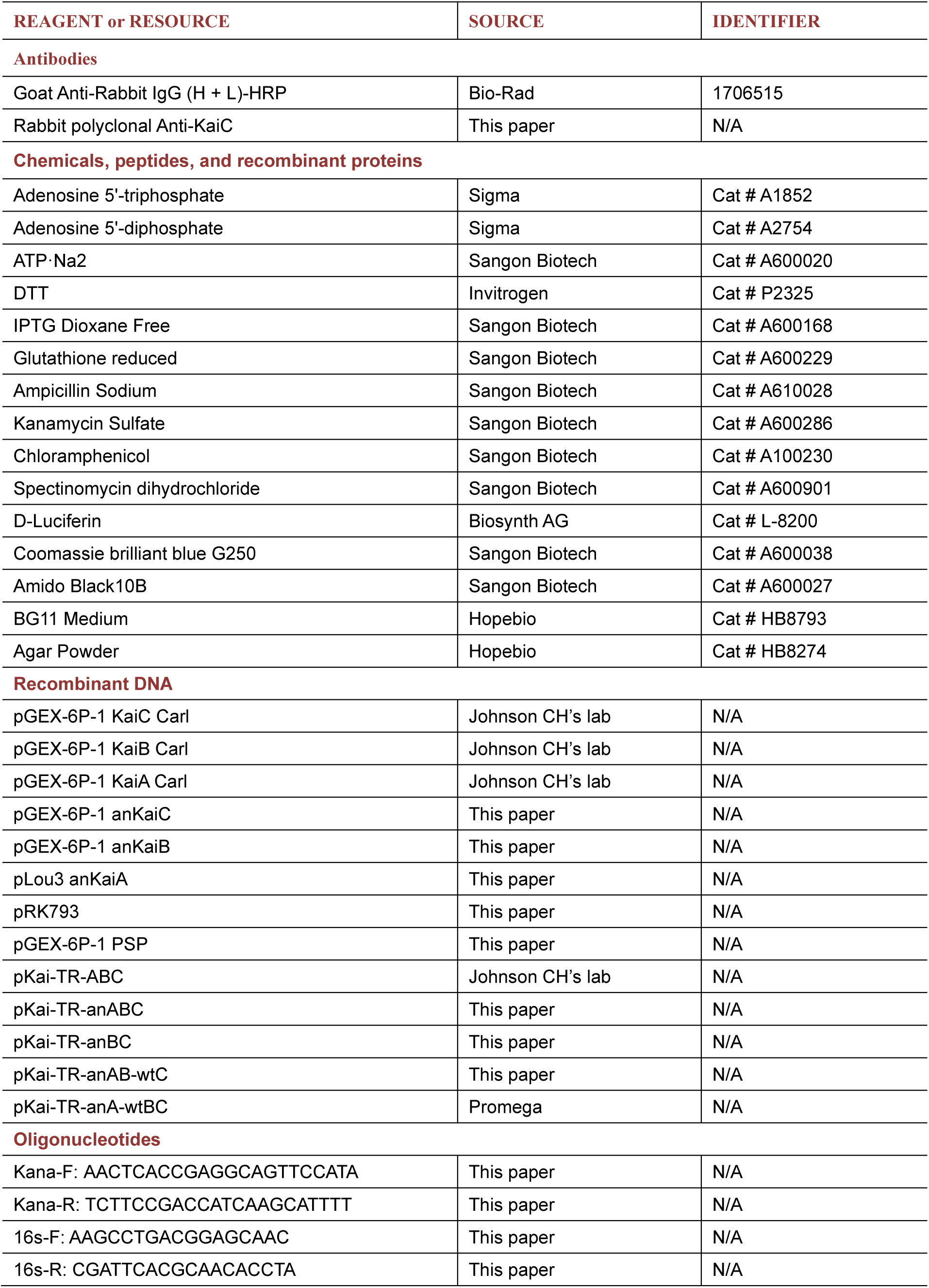
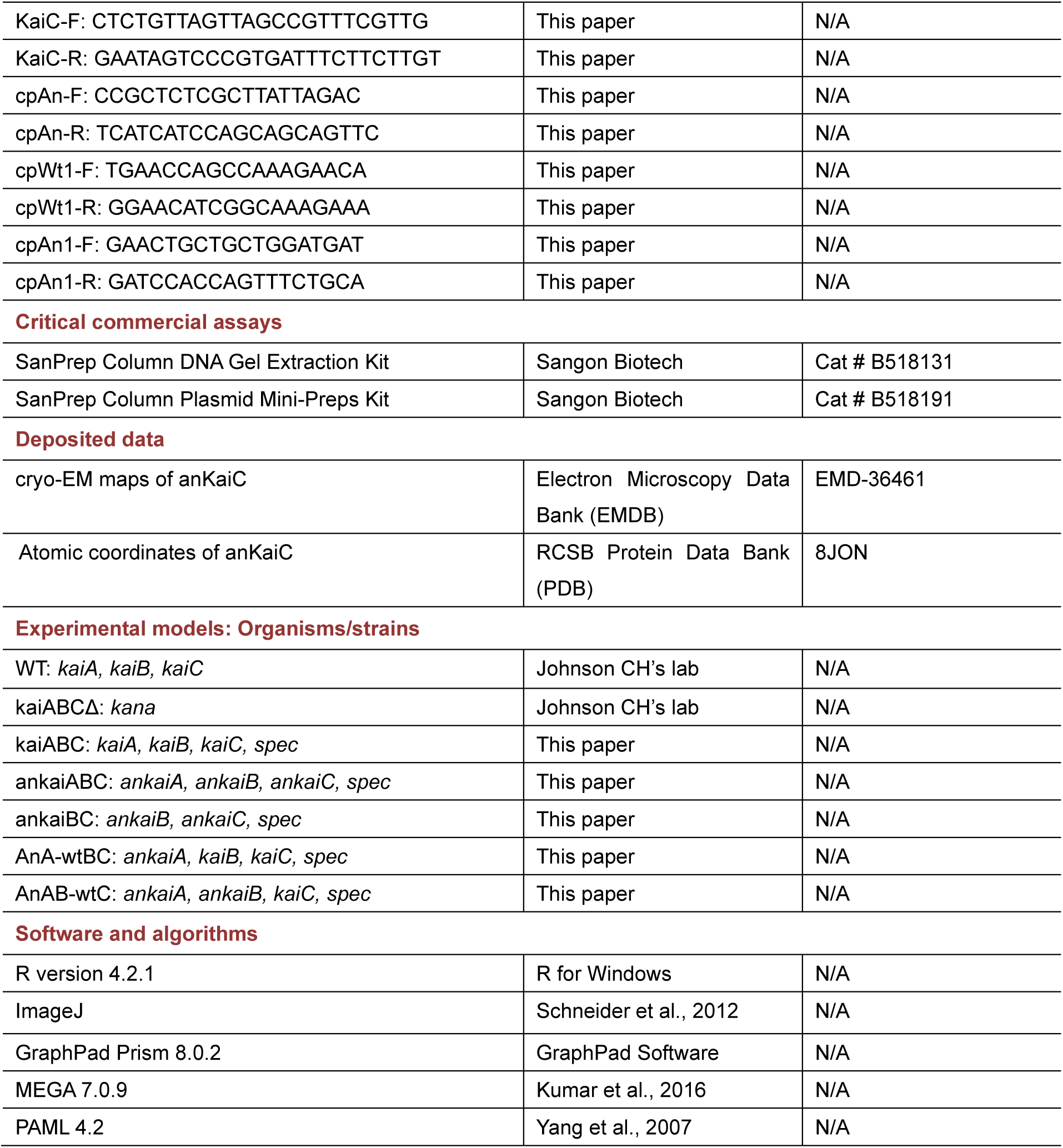

## RESOUCE AVAILABILITY

### Lead contact

Requests for strains or information should be directed to the lead contact, Jinhu Guo.

### Materials availability

Strains generated in this study are available upon request to the lead contact.

### Data and code availability

- Quantitative data sets are provided as supplemental source files with this publication. The cryo-EM maps of anKaiC have been deposited in the Electron Microscopy Data Bank (EMDB) under the accession code EMD-36461. The atomic coordinates have been deposited in the RCSB Protein Data Bank (PDB) under the accession code 8JON.
- This paper does not report original code.
- Any additional information required to reanalyze the data reported in this paper is available from the lead contact upon request.

## EXPERIMENTAL MODEL AND STUDY PARTICIPANT DETAILS

The *Synechococcus sp*. PCC7942 strain harbouring a *luxAB* reporter under the control of the *kaiBC* promoter to a neutral site, and the *Synechococcus sp*. PCC7942 *kaiABC*^Δ^ strain in which the locus of *KaiABC* is replaced by the kanamycin resistance gene (*kmr*)^49^ were gifts from Carl H. Johnson lab (Vanderbilt University, USA). To prepare strains expressing *anKaiABC* genes, *anKaiABC* genes were cloned and inserted into the *pKai-TR* plasmid which contains homologous flanking arms (526 nt upstream of *KaiA* and 288 nt downstream *KaiC*). The plasmids of *pKai-TR*-*KaiABC* and *pKai-TR*-*anKaiABC* were transformed into *kaiABC*^Δ^ and recombined into the locus of *kmr* using the natural transformation method. ^50–52^ Recombination of *KaiABC/anKaiABC* genes and *KaiABC* genes were also cloned and transformed in the same way. In the transformed strains, *KaiA*/*ankaiA* was driven by the *kaiA* promoter and *KaiBC*/*ankaiBC* were driven by the *kaiBC* promoter, respectively. Cyanobacteria was cultured at 30^°^C under constant light with an intensity of ∼50 μE m^−2^ s^−1^ in a light and temperature controllable incubator (Percival, USA). ^6^

## METHOD DETAILS

### Phylogenetic analysis of the ancestral sequences of cyanobacterial clock genes

The phylogenetic trees were constructed according to the 16S rRNA sequences and the divergence times along the cyanobacterial phylogeny were estimated under Bayesian relaxed molecular clocks. ^53,54^ The amino acid (AA) sequences of KaiA, KaiB, and KaiC from the *Cyanobacteria* AC group, ranging from Prochlorococcus marinus to *Synechococcussp* and *Synechococcus elongatus* PCC 7942, were used for ancestral sequence reconstruction. ^54^ These sequences were aligned with the program MUSCLE. The PAML package was used to infer the posterior AA probability per site under the LG model of protein evolution, with the phylogenetic tree constructed based on the 16S rRNA sequences. ^55^ We reconstructed the AA sequences of the ancestral KaiA, KaiB, and KaiC for the node at 0.95 Ga ago (Figure S1).

### Prokaryotic protein expression and purification

KaiABC gene cassettes were obtained from PCR amplification using the genomic DNA of *Synechococcus sp*. PCC7942 as templates, and anKaiABC genes were synthesized by Shenggong Co. Ltd. (Shanghai, China). Synthesis of the ancestral KaiA, KaiB, and KaiC were based on the *E. coli* codon usage table. The cassettes of KaiABC and anKaiBC genes were cloned and inserted into the *pGEX-6p-1* vector, and anKaiA was cloned into the *plou3* vector. The plasmids were transformed into *E. coli* BL21 and the target proteins were purified as reported previously. ^20^ For each protein, 1 L of culture was grown at 37 ^°^C to an OD600 of approximately 0.6, at which point it was cooled to 25^°^C and induced overnight with 100 μM IPTG. The cells were subsequently harvested, pelleted, and frozen at -80^°^C. anKaiABC and kaiABC proteins were purified with a GSTrap column (GE Healthcare, USA), Ni-NTA column (Qiagen, Germany) and Resource Q anion exchange column (GE Healthcare, USA). The protein concentration was determined by the Bradford method. The purity of anKaiA was > 90% as determined by SDS/PAGE. anKaiA was flash-frozen with liquid nitrogen and stored at -80°C.

### Cryo-EM sample preparation, data acquisition and processing, and model building

Three microliters of freshly purified anKaiC were applied to a glow discharged R1.2/1.3 holey copper grid (Quantifoil Micro Tools GmbH, USA). The grid was then blotted and plunged in precooled liquid ethane using a Vitrobot Mark IV (Thermo Fisher Scientific, USA). The cryo-EM data of anKaiC were collected under a Titan Krios G4 300 kV electron microscope (Thermo Fisher Scientific) equipped with a Falcon4 camera (Thermo Fisher Scientific, USA). Data collection was performed using *EPU* software (Thermo Fisher Scientific, USA) with defocus values ranging from 1.0 μm to 3.0 μm. Video frames were recorded in electron-event representation (EER) mode under a dose rate of 16.2 e^‒^/Å^2^/s, giving a total dose of 50 e^‒^/Å^2^. The nominal magnification was 165,000×, giving a calibrated pixel size of 0.71 Å. Beam-induced motion correction and dose weighting were performed using Relion3. ^56^ Contrast transfer function parameters were estimated using CTFFIND4. ^57^ Protein particles were automatically picked by the BoxNet convolutional neural network in the program Warp. ^58^ Rounds of reference-free 2D classification and 3D classification were performed using Relion3 to select clear and homogeneous particles. Higher-order aberrations, anisotropic magnification, and the x defocus value of each particle were refined, and the particles were polished. Then, the polished particles were 3D refined and generated the final reconstruction using Relion3. The resolution was generated using the gold standard FSC = 0.143 criterion. ^59,60^ The maps were sharpened using Phenix. ^61^ The atomic model of anKaiC was initially built by AlphaFold2^62^ and then manually tuned in COOT^63^ based on the electron density map. The model was refined and validated in Phenix. Refinement and validation statistics are summarized in Table S2. All Cryo-EM structure figures were prepared in UCSF ChimeraX. ^64^

### In vitro measurement of the autokinase and autophosphatase activities of KaiC/anKaiC

KaiC function analysis was performed as described with some modifications. ^20^ To assess its autophosphatase activity, initially anKaiC protein (0.2 μg/μL) was incubated in reaction buffer (50 mM Tris-HCl, pH 8.0, 150 mM NaCl, 1 mM DTT, 5 mM MgCl2, 1 mM ATP) at 30^°^C. Aliquots (5 µl) of the reaction mixtures were collected every 2 h and stored at -20°C. To assess the autokinase activity of anKaiC, after 15 h the reaction mixture was moved to 0^°^C and aliquots were sampled until 27 h after initiation. ^65^ All aliquots were resolved by electrophoresis on 10% separation gel at 175 V for 80 min. The gel was dyed with Coomassie brilliant blue G-250. NIH ImageJ software (version 1.51) was used for densitometric analysis. ^20^

### In vitro measurement of KaiA/anKaiA functions in promoting KaiC phosphorylation

KaiA functions to promote KaiC phosphorylation. ^20,66^ Initially, the KaiC/anKaic proteins were incubated in the solution with 0.2 μg/μL KaiC, 1 mM ATP, 5 mM MgCl2, 150 mM NaCl, 1 mM DTT and 50 mM Tris-HCl buffer (pH 8.0) at 30^°^C, to induce hypophosphorylation of KaiC/anKaiC. After 15 h of incubation, KaiA or anKaiA proteins were added to the system (final concentration: 0.05 μg/μL) and incubated for another 12 h.

Five microliter aliquots were taken out from each system every 3 h. All aliquots were resolved by electrophoresis on a 10% separation gel at 175 V for 80 min. The gel was dyed with Coomassie brilliant blue G-250. NIH ImageJ software (version 1.51) was used for densitometric analysis. ^20^

### In vitro measurement of KaiB/anKaiB functions in repressing KaiC phosphorylation

KaiB is capable of binding to KaiC which sequesters the KaiA-KaiC interaction and represses KaiC phosphorylation. ^67^ Initially, KaiA/anKaiA (final concentration: 0.05 μg/μL) and KaiC/anKaiC (0.2 μg/μL KaiC) were added to the solution with 1 mM ATP, 5 mM MgCl2, 150 mM NaCl, 1 mM DTT and 50 mM Tris-HCl buffer (pH 8.0), and incubated at 30°C for 12 h to induce hyperphosphorylation of KaiC or anKaiC. KaiB/anKaiB proteins were added to the solution (final concentration: 0.05 μg/μL) and incubated for 15 h. Five microliter aliquots were taken out from each system every 3 h. All aliquots were resolved by electrophoresis on a 10% separation gel at 175 V for 80 min. The gel was dyed with Coomassie brilliant blue G-250. NIH ImageJ software (version 1.51) was used for densitometric analysis. ^20^

### In vitro ATPase assay of KaiC

The ATPase activity of purified KaiC was measured by using an HPLC system. Briefly, the purified KaiC protein (0.5 pmol hexamer/µL) was incubated at 25^°^C in buffer [20 mM Tris·HCl (pH 8.0), 150 mM NaCl, and 5 mM MgCl2]. KaiC/anKaiC proteins were incubated with 1 mM ATP for the periods indicated in the presence (closed, solid line) or absence (open, dashed line) of KaiA (3.0 pmol/µL). The HPLC system (1290 Infinity, Agilent Technology, America) was equipped with a DAD detector; and a Perfect T3 (4.6X250 mm, 5 μm) column (Micropulite, China) as described. ATP and ADP in the reaction mixture were separated on the column at 40^°^C at a 1.0 ml/min flow rate. The detection wavelength was set at 260 nm. The mobile phase was composed of 50 mM NaH2PO4 buffer (pH 7.0) and 50% (vol/vol) acetonitrile. The ATPase activity was evaluated as a function of the amount of ADP produced.^37^

### In vitro reconstitution of KaiC phosphorylation oscillation

The detailed reconstitution of KaiC phosphorylation rhythm in vitro *was* performed as previously described. ^23,68^ Briefly, KaiC (0.2 µg/µl) was incubated with KaiA (0.05 µg/µl) and KaiB (0.05 µg/µl) in reaction buffer (50 mM Tris-HCl, pH 8.0, 150 mM NaCl, 1 mM DTT, 5 mM MgCl2, 1 mM ATP) at 30°C. Aliquots (5 µl) of the reaction mixtures were collected every 2 h, and the reaction was stopped by the addition of SDS loading buffer. Samples were subjected to SDS-PAGE on 10% gels followed by Coomassie Brilliant Blue staining. NIH ImageJ software (version 1.51) was used for densitometric analysis.

### Generation of KaiC multi-clone antibody and western blot analysis

KaiC protein was expressed in *E. coli*, extracted from inclusion bodies, and purified with a column containing Ni-NTA agarose (Qiagen, Germany). The resultant purity of KaiC protein was > 90%, which was used to immunize a rabbit. The serum containing antibodies against KaiC was extracted and validated. *Synechococcus* cells were resuspended in 100-200 μL of lysis buffer (25 mM Tris pH 8.0, 0.5 mM EDTA, 1 mM DTT and protease inhibitor cocktail), total protein was isolated and western blot experiments were performed as described previously. ^68^

### Transformation of cyanobacteria

Transformation of *cyanobacteria* was performed by following the previously published methods. ^45,46,63^ Briefly, 5 ml ∼ 10 ml of cyanobacteria cells were centrifuged at 1620 g for 5 min and washed in 10 ml of 10 mM NaCl. The cells were resuspended in 0.3 ml of BG11 medium. For transformation, 1 µl of miniprep plasmid (0.1 µg ∼ 0.3 µg) was added to the cells and incubated at 30^°^C overnight with gentle shaking in darkness. The transformed cells were grown on BG11-agar solid medium containing antibiotics, including chloramphenicol and spectinomycin. The transformed cells were grown on plates at 30^°^C under LL conditions until the formation of the transformed colonies. Single colonies were picked and grown in BG11 liquid media under LL. When the culture became green, PCR and sequencing were conducted for validation.

### Bioluminescence monitoring and associated analysis

*S. elongatus* strains expressing *PkaiBC-luxAB* were grown on petri dishes with BG11 media at 30°C. The strains were inoculated onto BG-11 solid medium in 90-mm plates and the bioluminescence intensity of the colonies was monitored by a CCD camera (iKon-M 934, Andor, Northern Ireland). The bacterial colonies were approximately 0.5 mm ∼ 1.0 mm in diameter and contained 100 - 1000 cells. Prior to monitoring, the plates were exposed to DD for 12 h to synchronize the circadian rhythms. Circadian rhythms of LUC activity were captured using a back-illuminated CCD sensor from e2v (CCD47-40) and normalized to the mean value over the time series. Fast Fourier transform-nonlinear least squares analysis of circadian parameters was conducted on a data window of ZT24-120. The bioluminescence activity of LUC fusion proteins was measured on a Packard TopCount^TM^ luminometer and used as a read-out of the state in LD12:12 and LL conditions, and the light intensity was 50 μE m^−2^ s^−1^ during the light regimes. N-decanal (3% v/v, dissolved in canola oil) was used as the substrate of luciferase and the final concentration was 3% v/v. ^69^ The bioluminescence data were analysed with R-4.2.1).

### Transmission electron microscopy (TEM) sample preparation and results analysis

*Synechococcus* cells were harvested by centrifugation at 1500 g, fixed in 2.5% glutaraldehyde and 2% paraformaldehyde prepared in 0.1 M PBS (pH 7.2) for 4 h at 4^°^C, and treated with 2% osmium tetroxide at room temperature overnight. Thereafter, the samples were gradient dehydrated and embedded in Spurr resin (Sigma-Aldrich Co. Ltd., USA). Ultrathin sections were cut and stained with aqueous uranyl acetate and lead citrate.^70^ The grids were examined with a JEM1400Flash electron microscope (JOEL Co. Ltd., Japan) operating at an acceleration voltage of 120 kV. For each replicate, the phenotypes of 100 randomly selected cells were observed and counted.

### Competition experiment with *Cyanobacteria* strains

The competition experiments were conducted as previously described except the determination of the composition of the mixture. ^6,70^ In the competition experiments, cyanobacteria were cultured on BG-11 solid or liquid medium under the indicated conditions with fluorescence lamps operated at 90-100 μE m^−2^ s^−1^. To calculate the ratios of different strains in the culture mixture, 50 µL of the mixture was taken out after the culture and grown on plates to obtain single colonies. Genomic DNA samples were extracted from > 100 single colonies and subjected to PCR and electrophoresis for genotyping to determine the composition of populations under competition. The sequences of the PCR primers are listed in Extended Data Table 1.

### Statistics

Statistical analyses were generally performed using R version 4.2.1 or with the software GraphPad Prism 8.0.2. The TEM results were analysed with unpaired *t*-test, and others analysed with two-tailed unpaired Student’s *t*-test. Data are plotted as mean ± standard deviation (s.d.) or mean ± standard error (s.e.m.) as indicated. * *P* < 0.05, ** *P* < 0.01, *** *P* < 0.001.

## REFERENCES

1. Bell-Pedersen, D., Cassone, V.M., Earnest, D.J., Golden, S.S., Hardin, P.E., Thomas, T.L., and Zoran, M.J. (2005). Circadian rhythms from multiple oscillators: lessons from diverse organisms. Nat. Rev. Genet. 6, 544–556.

2. Yerushalmi, S., and Green, R.M. (2009). Evidence for the adaptive significance of circadian rhythms. Ecol. Lett. 12, 970–981.

3. Johnson, C.H., Zhao, C., Xu, Y., and Mori, T. (2017). Timing the day: what makes bacterial clocks tick? Nat. Rev. Microbiol. 15, 232–242.

4. Eelderink-Chen, Z., Bosman, J., Sartor, F., Dodd, A.N., Kovács, Á.T., and Merrow, M. (2021). A circadian clock in a nonphotosynthetic prokaryote. Sci. Adv. 7, eabe2086.

5. Grobbelaar, N., Huang, T.C., Lin, H.Y., and Chow, T.J. (1986). Dinitrogen-fixing endogenous rhythm in Synechococcus RF-1. FEMS Microbiol. Lett. 37, 173–177.

6. Ouyang, Y., Andersson, C.R., Kondo, T., Golden, S.S., and Johnson, C.H. (1998). Resonating circadian clocks enhance fitness in cyanobacteria. Proc. Natl. Acad. Sci. USA 95, 8660–8664.

7. Woelfle, M.A., Ouyang, Y., Phanvijhitsiri, K., and Johnson, C.H. (2004). The adaptive value of circadian clocks: an experimental assessment in cyanobacteria. Curr. Biol. 14,1481–1486.

8. Dodd, A.N., Salathia, N., Hall, A., Kévei, E., Tóth, R., Nagy, F., Hibberd, J.M., Millar, A.J., and Alex, AR Webb. (2005). Plant circadian clocks increase photosynthesis, growth, survival, and competitive advantage. Science 309, 630–633.

9. Highkin, H.R., and Hanson, J.B. (1954). Possible Interaction between Light-dark Cycles and Endogenous Daily Rhythms on the Growth of Tomato Plants. Plant Physiol. 29, 301–302.

10. Mwimba, M., Karapetyan, S., Liu, L., Marqués, J., McGinnis, E.M., Buchler, N.E., and Dong, X. (2018). Daily humidity oscillation regulates the circadian clock to influence plant physiology. Nat. Commun. 9, 4290.

11. Ma, H., Li, L., Yan, J., Zhang, Y., Ma, X., Li, Y., Yuan, Y., Yang, X., Yang, L., and Guo, J. (2021). The Resonance and Adaptation of Neurospora crassa Circadian and Conidiation Rhythms to Short Light-Dark Cycles. J. Fungi 8, 27.

12. Schopf, J.W., and Packer, B.M. (1987). Early Archean (3.3-billion to 3.5-billion-year-old) microfossils from Warrawoona Group, Australia. Science 237, 70–73.

13. Ishiura, M., Kutsuna, S., Aoki, S., Iwasaki, H., Andersson, C.R., Tanabe, A., Golden, S.S., Johnson. C.H., and Kondo, T. (1998). Expression of a gene cluster kaiABC as a circadian feedback process in cyanobacteria. Science 281, 1519–1523.

14. Johnson, C.H., Golden, S.S., and Kondo, T. (1998). Adaptive significance of circadian programs in cyanobacteria. Trends. Microbiol. 6, 407–410.

15. Nishiwaki, T., Satomi, Y., Nakajima, M., Lee, C., Kiyohara, R., Kageyama, H., Kitayama, Y., Temamoto, M., Yamaguchi, A., Hijikata, A., et al. (2004). Role of KaiC phosphorylation in the circadian clock system of Synechococcus elongatus PCC 7942. Proc. Natl. Acad. Sci. USA 101, 13927–13932.

16. Hayashi, F., Itoh, N., Uzumaki, T., Iwase, R., Tsuchiya, Y., Yamakawa, H., Morishita, M., Onai, K., Itoh, S., and Ishiura, M. (2004). Roles of two ATPase-motif-containing domains in cyanobacterial circadian clock protein KaiC. J. Biol. Chem. 279, 52331–52337.

17. Swan, J.A., Sandate, C.R., Chavan, A.G., Freeberg, A.M., Etwaru, D., Ernst, D.C., Palacios, J.G., Golden, S.S., LiWang, A., Lander, G.C., et al. (2022). Coupling of distant ATPase domains in the circadian clock protein KaiC. Nat. Struct. Mol. Biol. 29, 759–766.

18. Kim, Y.I., Dong, G., Carruthers, C.W. Jr, Golden, S.S., and LiWang, A. (2008). The day/night switch in KaiC, a central oscillator component of the circadian clock of cyanobacteria. Proc. Natl. Acad. Sci. USA 105, 12825–12830.

19. Nakajima, M., Imai, K., Ito, H., Nishiwaki, T., Murayama, Y., Iwasaki, H., Oyama, T., and Kondo, T. (2005). Reconstitution of circadian oscillation of cyanobacterial KaiC phosphorylation in vitro. Science 308, 414–415.

20. Dvornyk, V. (2006). Evolution of the circadian clock mechanism in prokaryotes. Isr. J. Ecol. Evol. 52, 343–357.

21. Baca, I., Sprockett, D., and Dvornyk, V. (2010). Circadian input kinases and their homologs in cyanobacteria: evolutionary constraints versus architectural diversification. J. Mol. Evol. 70, 453–465.

22. Holtzendorff, J., Partensky, F., Mella, D., Lennon, J.-F., Hess, W.R., and Garczarek, L. (2008). Genome streamlining results in loss of robustness of the circadian clock in the marine cyanobacterium Prochlorococcus marinus PCC 9511. J. Biol. Rhythms. 23, 187–199.

23. Axmann, I.M., Dühring, U., Seeliger, L., Arnold, A., Vanselow, J.T., Kramer, A., and Wilde, A. (2009). Biochemical evidence for a timing mechanism in prochlorococcus. J. Bacteriol. 191, 5342–5347.

24. Ma, P., Mori, T., Zhao, C., Thiel, T., and Johnson, C.H. (2016). Evolution of KaiC-Dependent Timekeepers: A Proto-circadian Timing Mechanism Confers Adaptive Fitness in the Purple Bacterium Rhodopseudomonas palustris. PLoS Genet. 12, e1005922.

25. Min, H., Guo, H., and Xiong, J. (2005). Rhythmic gene expression in a purple photosynthetic bacterium, Rhodobacter sphaeroides. FEBS Lett. 579, 808–812.

26. Scrutton, C.T., and Hipkin, R.G. (1973). Long-term changes in the rotation rate of the Earth. Earth-Sci. Rev. 9, 259–274.

27. Sonett, C.P., Kvale, E.P., Zakharian, A., Chan, M.A., and Demko, T.M. (1996). Late Proterozoic and Paleozoic Tides, Retreat of the Moon, and Rotation of the Earth. Science 273, 100–104.

28. Spalding, C., and Fischera, W.W. (2019). A shorter Archean day-length biases interpretations of the early Earth’s climate. Earth Planet Sci. Lett. 514, 28–36.

29. Ditty, J.L., Mackey, S.R., and Johnson, C.H. (2009). Bacterial circadian programs (Springer-Verlag Berlin Heidelberg); pp. 241–258.

30. Zhao, Z., Zhou, Y., and Ji, G. (2007). The periodic growth increments of biological shells and the orbital parameters of Earth-Moon system. Environ. Geol. 51, 1271–1277.

31. Gaucher, E.A., Thomson, J.M., Burgan, M.F., and Benner, S.A. (2003). Inferring the palaeoenvironment of ancient bacteria on the basis of resurrected proteins. Nature 425, 285–288.

32. Hochberg, G.K.A., and Thornton, J.W. (2017). Reconstructing Ancient Proteins to Understand the Causes of Structure and Function. Annu. Rev. Biophys. 46, 247–269.

33. Dvornyk, V., Vinogradova, O., and Nevo, E. (2003). Origin and evolution of circadian clock genes in prokaryotes. Proc. Natl Acad. Sci. USA 100, 2495–2500.

34. Egli, M., Pattanayek, R., Sheehan, J.H., Xu, Y., Mori, T., Smith, J.A., and Johnson, C.H. (2013). Loop-loop interactions regulate KaiA-stimulated KaiC phosphorylation in the cyanobacterial KaiABC circadian clock. Biochemistry 52, 1208–1220.

35. Tseng, R., Chang, Y.G., Bravo, I., Latham, R., Chaudhary, A., Kuo, N.W., and Liwang, A. (2014). Cooperative KaiA-KaiB-KaiC interactions affect KaiB/SasA competition in the circadian clock of cyanobacteria. J. Mol. Biol. 426, 389–402.

36. Terauchi, K., Kitayama, Y., Nishiwaki, T., Miwa, K., Murayama, Y., Oyama, T., and Kondo, T. (2007). ATPase activity of KaiC determines the basic timing for circadian clock of cyanobacteria. Proc. Natl. Acad. Sci. USA 104, 16377–16381.

37. Abe, J., Hiyama, T.B., Mukaiyama, A., Son, S., Mori, T., Saito, S., Osako, M., Wolanin, J., Yamashita, E., Kondo, T., et al. (2015). Circadian rhythms. Atomic-scale origins of slowness in the cyanobacterial circadian clock. Science 349, 312–316.

38. Pattanayek, R., Mori, T., Xu, Y., Pattanayek, S., Johnson, C.H., and Egli, M. (2009). Structures of KaiC circadian clock mutant proteins: a new phosphorylation site at T426 and mechanisms of kinase, ATPase and phosphatase. PLoS One 4, e7529.

39. Nobel, P.S. (2020). 5 - Photochemistry of Photosynthesis. Chapter in Physiochem Environ Plant Physiol (5rd Edition) (Academic Press); pp. 259–307.

40. Krasinsky, G.A. and Brumberg, V.A. (2004). Secular increase of astronomical unit from analysis of the major planet motions, and its interpretation. Celest. Mech. Dyn. Astron, 90, 267–288.

41. Chew, J., Leypunskiy, E., Lin, J., Murugan, A., and Rust, M.J. (2018). High protein copy number is required to suppress stochasticity in the cyanobacterial circadian clock. Nat. Commun. 9, 3004.

42. Cann, R.L., Stoneking, M., and Wilson, A.C. (1987). Mitochondrial DNA and human evolution. Nature 325, 31–36.

43. Kawamoto, N., Ito, H., Tokuda, I.T., and Iwasaki, H. (2020). Damped circadian oscillation in the absence of KaiA in Synechococcus. Nat. Commun. 11, 2242.

44. Nishiwaki, T., Iwasaki, H., Ishiura, M., and Kondo, T. (2000). Nucleotide binding and autophosphorylation of the clock protein KaiC as a circadian timing process of cyanobacteria. Proc. Natl. Acad. Sci. USA 97, 495–499.

45. Nishiwaki, T., and Kondo, T. (2012). Circadian autodephosphorylation of cyanobacterial clock protein KaiC occurs via formation of ATP as intermediate. J. Biol. Chem. 287, 18030–18035.

46. Ma, P., Mori, T., Zhao, C., Thiel, T., and Johnson CH. (2016). Evolution of KaiC-Dependent Timekeepers: A Proto-circadian Timing Mechanism Confers Adaptive Fitness in the Purple Bacterium Rhodopseudomonas palustris. PLoS Genet. 12, e1005922.

47. Pitsawong, W., Pádua, R.A.P., Grant, T., Hoemberger, M., Otten, R., Bradshaw, N., Grigorieff, N., and Kern, D. (2023). From primordial clocks to circadian oscillators. Nature 616, 183–189.

48. Poulton, S.W. (2017). Early phosphorus redigested. Nat. Geosci. 10, 75–76.

49. Nishimura, H., Nakahira, Y., Imai, K., Tsuruhara, A., Kondo, H., Hayashi, H., Hirai, M., Saito, H., and Kondo, T. (2002). Mutations in KaiA, a clock protein, extend the period of circadian rhythm in the cyanobacterium Synechococcus elongatus PCC 7942. Microbiology 148, 2903–2909.

50. Edgar, R.S., Green, E.W., Zhao, Y., van Ooijen, G., Olmedo, M., Qin, X., Xu, Y., Pan, M., Valekunja, U.K., Feeney, K.A., et al. (2012). Peroxiredoxins are conserved markers of circadian rhythms. Nature 485, 459–464.

51. Golden, S.S., Brusslan, J., and Haselkorn, R. (1987). Genetic engineering of the cyanobacterial chromosome. Methods Enzymol. 153, 215–231.

52. Vioque, A. (2007). Advances in Experimental Medicine and Biology, vol 616 (Springer).

53. Schirrmeister, B.E., Antonelli, A., and Bagheri, H.C. (2011). The origin of multicellularity in cyanobacteria. BMC Evol. Biol. 11, 45.

54. Schirrmeister, B.E., de Vos, J.M., Antonelli, A., and Bagheri, H.C. (2013). Evolution of multicellularity coincided with increased diversification of cyanobacteria and the Great Oxidation Event. Proc. Natl. Acad. Sci. USA 110, 1791–1796.

55. Yang, Z. (2007). PAML 4: phylogenetic analysis by maximum likelihood. Mol. Biol. Evol. 24, 1586–1591.

56. Zivanov, J., Nakane, T., Forsberg, B.O., Kimanius, D., Hagen, W.J., Lindahl, E., Scheres, S.H. (2018). New tools for automated high-resolution cryo-EM structure determination in RELION-3. Elife 7, e42166.

57. Rohou, A., and Grigorieff, N. (2015). CTFFIND4: Fast and accurate defocus estimation from electron micrographs. J. Struct. Biol. 192, 216–221.

58. Tegunov, D., and Cramer, P. (2019). Real-time cryo-electron microscopy data preprocessing with Warp. Nat. Methods 16, 1146–1152.

59. Henderson, R., Sali, A., Baker, M.L., Carragher, B., Devkota, B., Downing, K.H., Egelman, E.H., Feng, Z., Frank, J., and Grigorieff, N., et al. (2012). Outcome of the first electron microscopy validation task force meeting. Structure 20, 205–214.

60. Scheres, S.H., and Chen, S. (2020). Prevention of overfitting in cryo-EM structure determination. Nat. Methods. 9, 853–854.

61. Liebschner, D., Afonine, P.V., Baker, M.L., Bunkóczi, G., Chen, V.B., Croll, T.I., Hintze, B., Hung, L.W., Jain, S., and McCoy, A.J., et al. (2019). Macromolecular structure determination using X-rays, neutrons and electrons: recent developments in Phenix. Acta Crystallogr. D Struct. Biol. 75, 861–877.

62. Jumper, J., Evans, R., Pritzel, A., Green, T., Figurnov, M., Ronneberger, O., Tunyasuvunakool, K., Bates, R., Žídek, A., Potapenko, A., et al. (2021). Highly accurate protein structure prediction with AlphaFold. Nature 596, 583–589.

63. Emsley, P., Lohkamp, B., Scott, W.G., and Cowtan, K. (2010). Features and development of Coot. Acta Crystallogr. D Biol. Crystallogr. 66, 486–501.

64. Goddard, T.D., Huang, C.C., Meng, E.C., Pettersen, E.F., Couch, G.S., Morris, J.H., and Ferrin, T.E. (2018). UCSF ChimeraX: Meeting modern challenges in visualization and analysis. Protein Sci. 27, 14–25.

65. Snijder, J., Burnley, R.J., Wiegard, A., Melquiond, A.S., Bonvin, A.M., Axmann, I.M., and Heck, A.J. (2014). Insight into cyanobacterial circadian timing from structural details of the KaiB-KaiC interaction. Proc. Natl. Acad. Sci. USA 111, 1379–1384.

66. Iwasaki, H., Nishiwaki, T., Kitayama, Y., Nakajima, M., and Kondo, T. (2002). KaiA-stimulated KaiC phosphorylation in circadian timing loops in cyanobacteria. Proc. Natl. Acad. Sci. USA 99,15788–15793.

67. Kitayama, Y., Iwasaki, H., Nishiwaki, T., and Kondo, T. (2003). KaiB functions as an attenuator of KaiC phosphorylation in the cyanobacterial circadian clock system. EMBO J. 22, 2127–2134.

68. Kim, Y.I., Boyd, J.S., Espinosa, J., and Golden, S.S. (2015). Detecting KaiC phosphorylation rhythms of the cyanobacterial circadian oscillator in vitro and in vivo. Methods Enzymol. 551, 153–173.

69. Mackey, S.R., Ditty, J.L., Clerico, E.M., and Golden, S.S. (2007). Detection of rhythmic bioluminescence from luciferase reporters in cyanobacteria. Methods Mol. Biol. 362, 115–129.

70. Zhang, Q., Cui, M., Huang, X., Zheng, H., Huang, J., Fang, L., Li, K., and Zhang, J. (2004). The life cycle of SARS coronavirus in Vero E6 cells. J. Med. Virol. 73, 332–337.

