## Supplemental Figures for "Ancient cyanobacterial proto-circadian clock adapted to short day-night cycles ∼ 0.95 billion years ago"

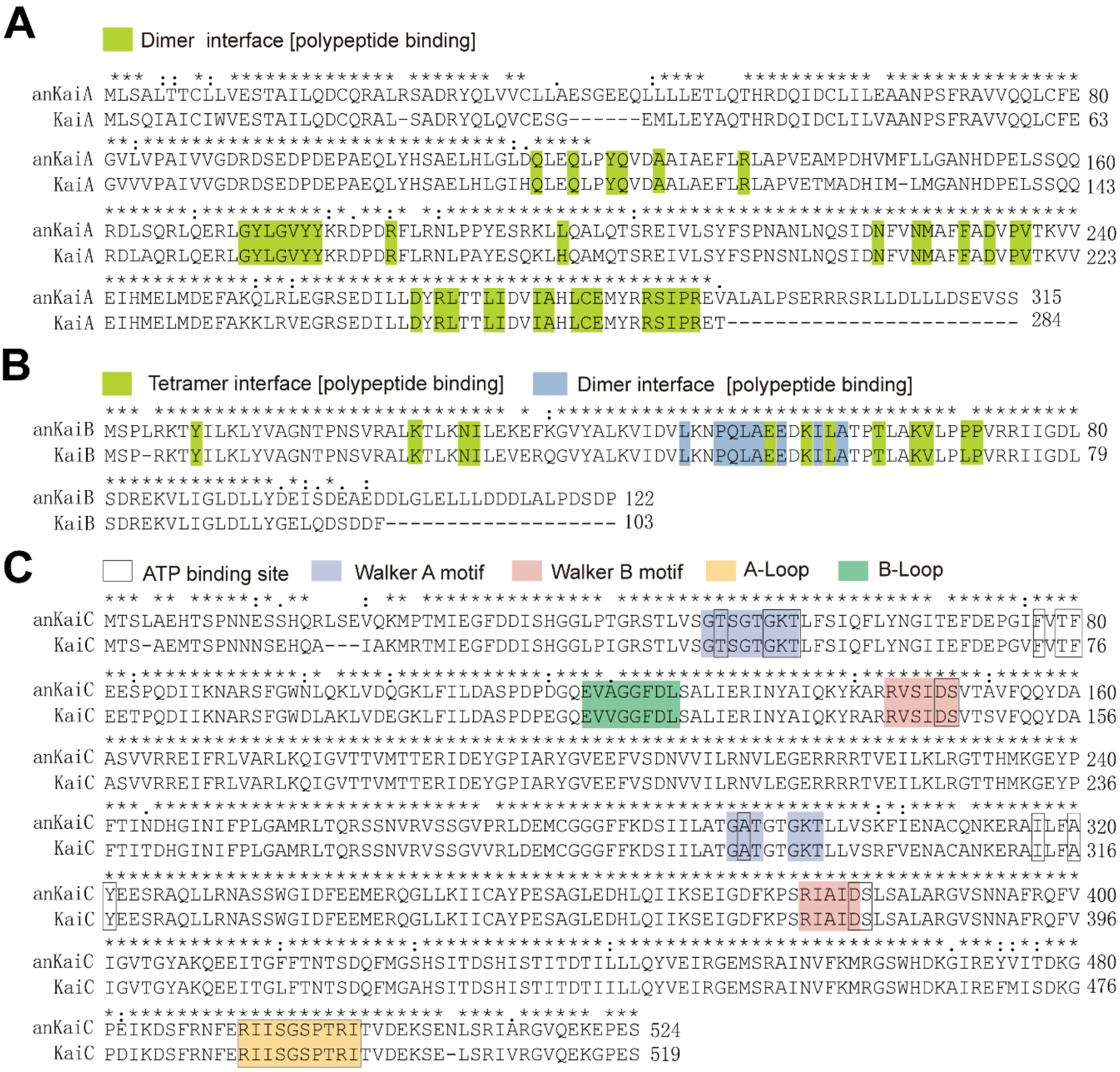


**Figure S1. Alignments of cyanobacterial KaiABC and anKaiABC sequences, related to Figure 1.**

(A to C) Protein sequence alignments between KaiA and anKai (A), KaiB and anKaiB (B), and KaiC and anKaiC (C) are shown. Critical domains are highlighted in different colours.

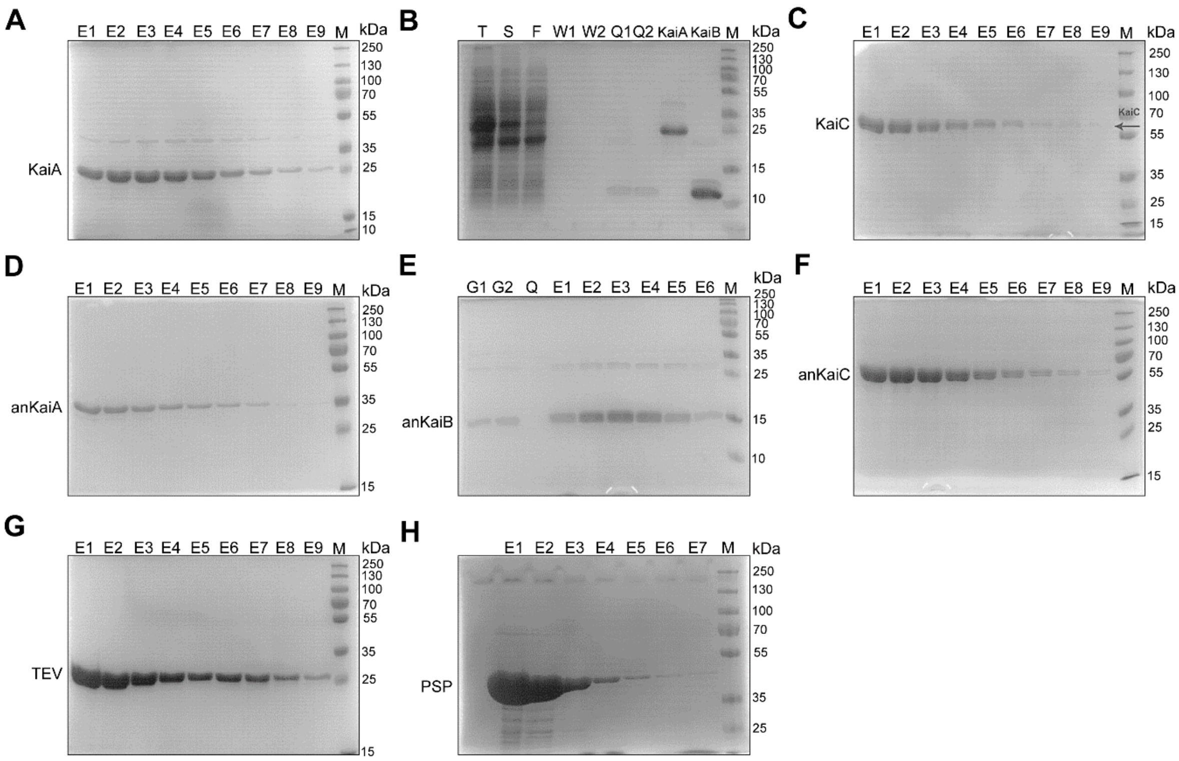


**Figure S2. SDS–polyacrylamide gel electrophoresis of purified prokaryotically expressed KaiABC and anKaiABC proteins, related to Figure 1.**

(A-C) SDS–polyacrylamide gel electrophoresis of purified KaiABC proteins.

(D-F) Electrophoresis of purified KaiABC proteins. In (E), the proteins were previously purified with GSTrap columns and here further purified with GSTrap columns once more.

(G,H) SDS–polyacrylamide gel electrophoresis of purified Tobacco Etch Virus protease (TEVp) (G), and PreScission Protease (PSP) (H) proteins. PSP was used to eliminate the MBP fusion tag and TEVp was used to remove the GST fusion tag.

In (A,C,D and F), Q columns were used for purification; in (B,E and H), GSTrap columns were used for purification; and in (G), Ni-NTA columns were used for protein purification. T: total cell lysates; S: supernatant; F: flow-through fractions; W1/2, fractions of wash 1/2; M, protein marker; E1-E9, elution fractions; Q1/2, proteins obtained by purification and following hyperfiltration concentration. All the purity values of these purified proteins were > 90%, which were measured and assessed with NIH ImageJ analysis software (version 1.51).

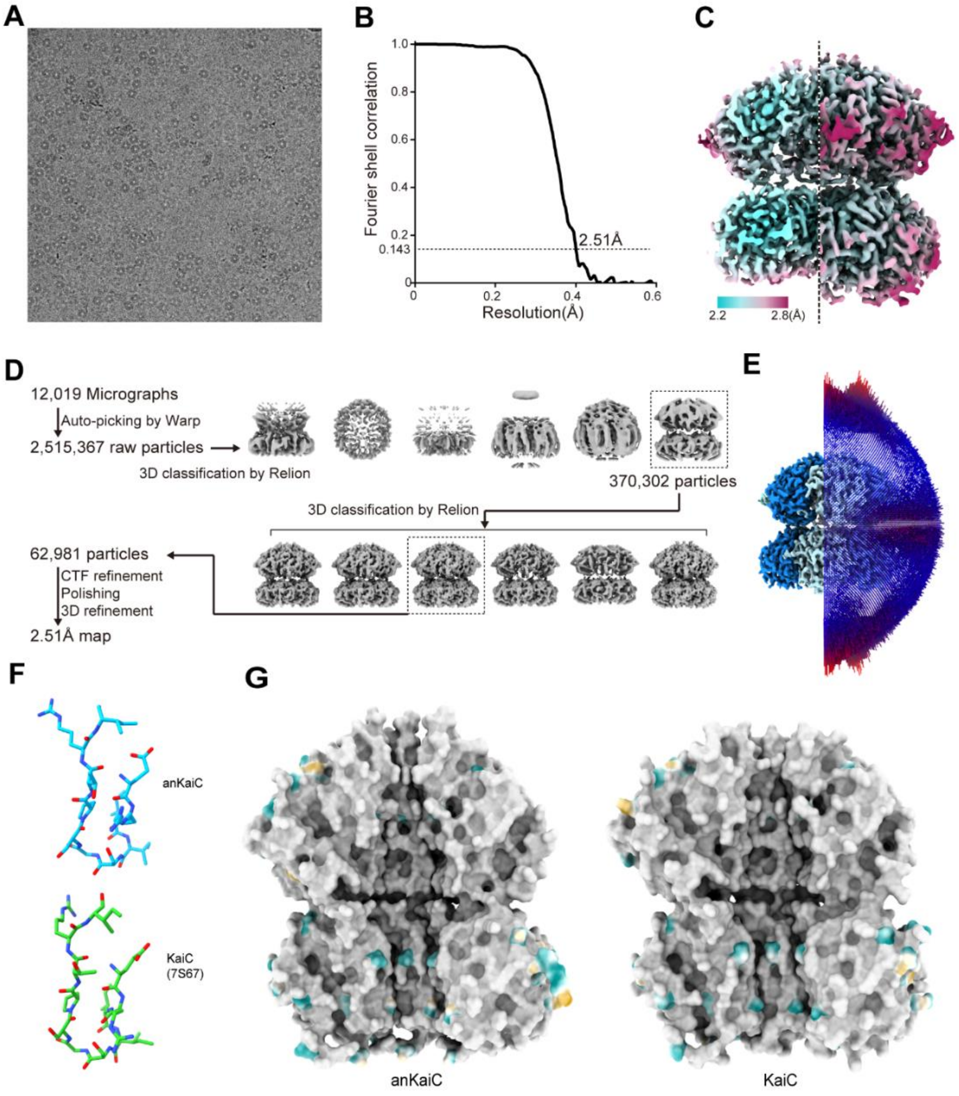


**Figure S3. Cryo-EM analysis of the anKaiC, related to Figure 1.**

(A) Representative cryo-EM micrograph of anKaiC.

(B) Fourier shell correlation curve of the anKaiC reconstruction.

(C) Local resolution map of anKaiC reconstruction.

(D) Image processing pipeline for anKaiC. Selected classes in each 3D classification are labelled by dotted line boxes.

(E) Angular distribution of particle orientations used in the final reconstruction.

(F) Conserved structure of partial A-loop between anKaiC (coloured blue) and KaiC (PDB: 3DVL7S67, coloured in pink green).

(G) The inside surface of anKaiC showing similarity to that of KaiC WT (PDB: 7S67). The surface (inside view) with identical amino acid residues in both anKaiC and KaiC are coloured in grey, the different amino acids between anKaiC and KaiC with hydrophilic residues are shown in cyan, and hydrophobic residues are shown in golden rod.

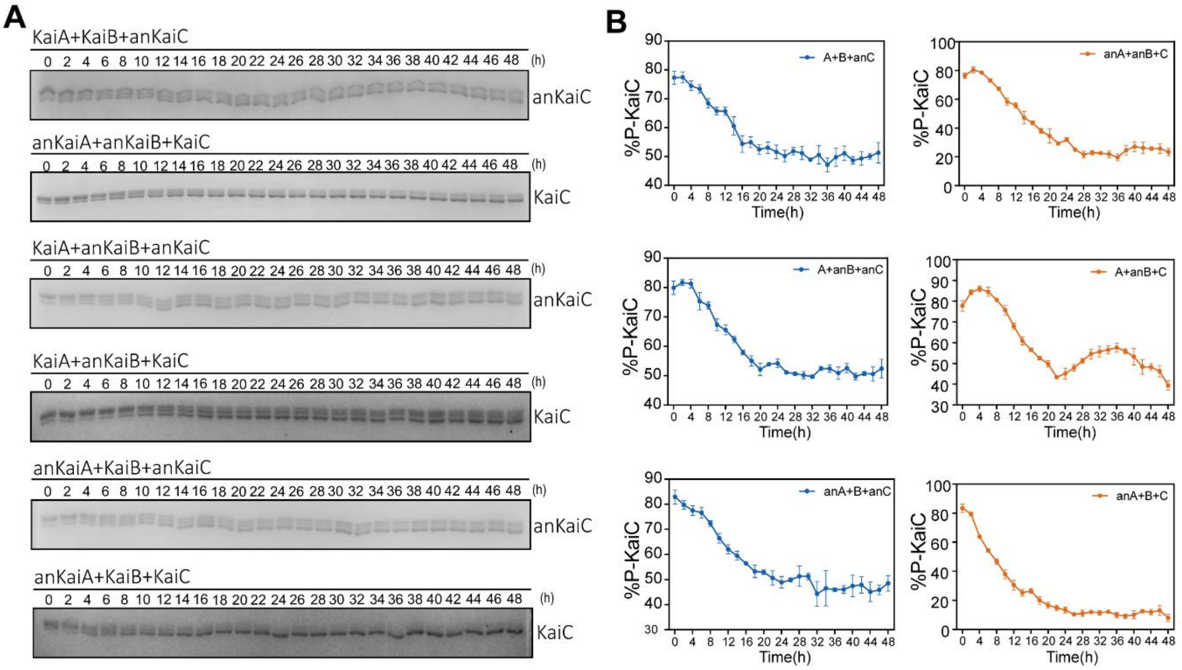


**Figure S4. Analysis of *in vitro* and *in vivo* rhythmicities in strains expressing different combinations of KaiABC proteins, related to Figure 3.**

(A) *In vitro* phosphorylation of anKaiC/KaiC proteins in different combinations. Representative results of Coomassie brilliant blue staining are shown.

(B) Statistical results of bioluminescence rhythms of the indicated combinations. The percentage of hyperphosphorylated KaiC was calculated. Data are means ± s.d. n=3.

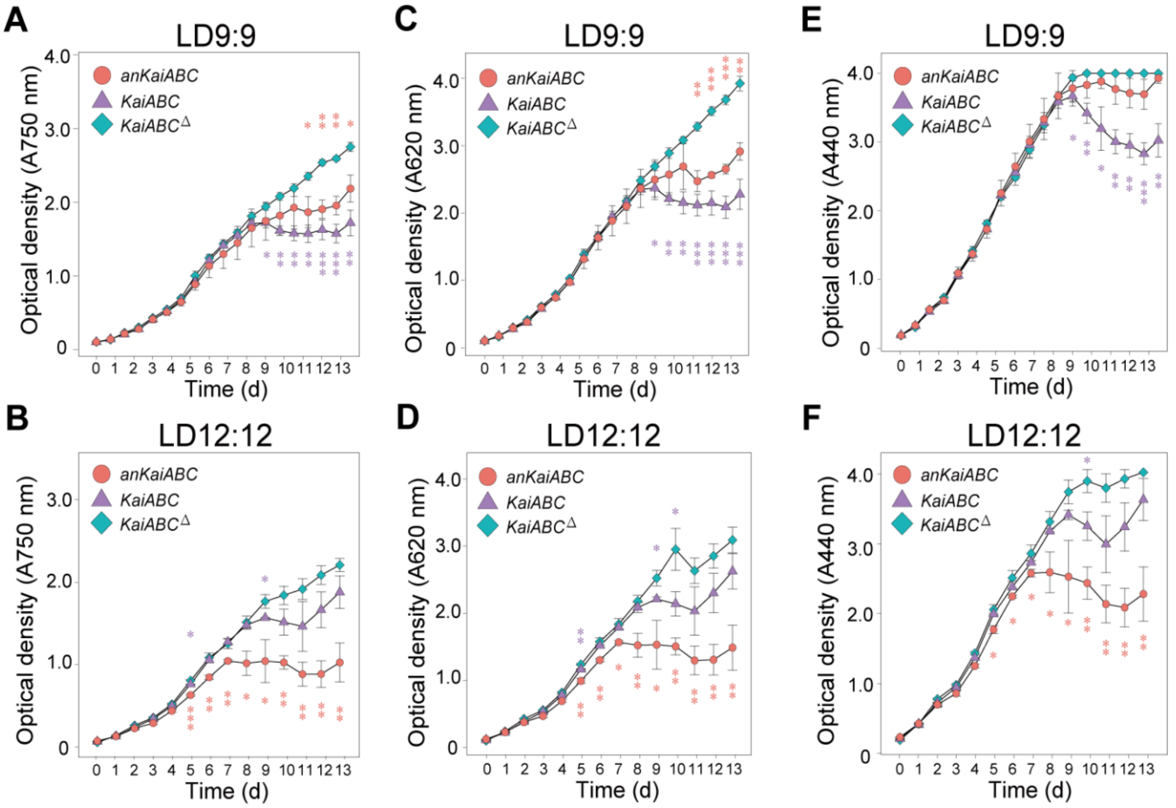


**Figure S5. Contribution of the ancient cyanobacterial circadian clock system to growth, related to Figure 4.**

(A,B) Growth curves of the indicated strains in LD9:9 and LD12:12.

(C,D) Phycobilin content measured by light absorption at 620 nm.

(E,F) Content of chlorophyll a measured by light absorption at 440 nm.

Data are means ± s.d. n=3 (A-F). Asterisks in purple denote significance between anKaiABC and KaiABC$^{\triangle}$, asterisks in red denote significance between KaiABC and KaiABC$^{\triangle}$.

**Table S1. Summary of model refinement and validation statistics of anKaiC.**

| | |
|---|---|
| **Composition** | |
| **Chains** | 6 |
| **Non-hydrogen atoms** | 45,942 |
| **Residues** | 2,910 |
| **Bonds (RMSD)** | |
| **Length** | 0.003 |
| **Angles** | 0.577 |
| **MolProbity score** | 1.36 |
| **Clash score** | 6.60 |
| **Ramachandran plot (%)** | |
| **Outliers** | 0.00 |
| **Allowed** | 1.86 |
| **Favored** | 98.14 |
| **Rotamer outliers (%)** | 0.96 |
| **Cβ outliers (%)** | 0.00 |
| **Model vs. Data** | |
| **CC (mask)** | 0.77 |
| **CC (volume)** | 0.70 |

**Table S2. PCR primers used in genotyping.**

| Gene | Usage | Primer name | Sequence (5’ - 3’) |
|---|---|---|---|
| *Kana* | genotyping | Kana-F | AACTCACCGAGGCAGTTCCATA |
| | | Kana-R | TCTTCCGACCATCAAGCATTTT |
| 16S ribosomal RNA | genotyping | 16s-F | AAGCCTGACGGAGCAAC |
| | | 16s-R | CGATTCACGCAACACCTA |
| *KaiC* | genotyping | KaiC-F | CTCTGTTAGTTAGCCGTTTCGTTG |
| | | KaiC-R | GAATAGTCCCGTGATTTCTTCTTGT |
| *anKaiA+anKaiB* | genotyping | cpAn-F | CCGCTCTCGCTTATTAGAC |
| | | cpAn-R | TCATCATCCAGCAGCAGTTC |
| *KaiA* | genotyping | cpWt1-F | TGAACCAGCCAAAGAACA |
| | | cpWt1-R | GGAACATCGGCAAAGAAA |
| *anKaiB+anKaiC* | genotyping | cpAn1-F | GAACTGCTGCTGGATGAT |
| | | cpAn1-R | GATCCACCAGTTTCTGCA |
